# Detection of long repeat expansions from PCR-free whole-genome sequence data

**DOI:** 10.1101/093831

**Authors:** Egor Dolzhenko, Joke J.F.A. van Vugt, Richard J. Shaw, Mitchell A. Bekritsky, Marka van Blitterswijk, Giuseppe Narzisi, Subramanian S. Ajay, Vani Rajan, Zoya Kingsbury, Sean J. Humphray, Raymond D. Schellevis, William J. Brands, Matt Baker, Rosa Rademakers, Maarten Kooyman, Gijs H.P. Tazelaar, Michael A. van Es, Russell McLaughlin, William Sproviero, Aleksey Shatunov, Ashley Jones, Ahmad Al Khleifat, Alan Pittman, Sarah Morgan, Orla Hardiman, Ammar Al-Chalabi, Chris Shaw, Bradley Smith, Edmund J. Neo, Karen Morrison, Pamela J. Shaw, Catherine Reeves, Lara Winterkorn, Nancy S. Wexler, The US-Venezuela Collaborative Research Group, David E. Housman, Christopher Ng, Alina Li, Ryan J. Taft, Leonard H. van den Berg, David R. Bentley, Jan H. Veldink, Michael A. Eberle

**Affiliations:** Illumina Inc., 5200 Illumina Way, San Diego, CA, USA; Department of Neurology, Brain Center Rudolf Magnus, University Medical Center Utrecht, Utrecht, The Netherlands; Illumina Cambridge Ltd., Chesterford Research Park, Little Chesterford, UK; Repositive Ltd., Future Business Centre, Kings Hedges Rd, Cambridge, UK; Department of Neuroscience, Mayo Clinic, Jacksonville, FL, USA; New York Genome Center, 101 Avenue of the Americas, New York, NY, USA; SURFsara, Science Park 140, Amsterdam, The Netherlands; Academic Unit of Neurology, Trinity College Dublin, Trinity Biomedical Sciences Institute, Dublin, Republic of Ireland; Department of Neurology, Beaumont Hospital, Dublin, Republic of Ireland; Department of Basic and Clinical Neuroscience, Maurice Wohl Clinical Neuroscience Institute, King’s College London, London, UK; Department of Molecular Neuroscience, UCL Institute of Neurology, London, UK; University of Southampton, Southampton, UK; Sheffield Institute for Translational Neuroscience, University of Sheffield, Sheffield, UK; Columbia University, 1051 Riverside Drive, New York, NY USA; Hereditary Disease Foundation, 3960 Broadway, 6^th^ floor, New York, NY, USA; The US-Venezuela Collaborative Research Group; Massachusetts Institute of Technology, 77 Massachusetts Ave., Cambridge, MA, USA

## Abstract

Identifying large repeat expansions such as those that cause amyotrophic lateral sclerosis (ALS) and Fragile X syndrome is challenging for short-read (100-150 bp) whole genome sequencing (WGS) data. A solution to this problem is an important step towards integrating WGS into precision medicine. We have developed a software tool called ExpansionHunter that, using PCR-free WGS short-read data, can genotype repeats at the locus of interest, even if the expanded repeat is larger than the read length. We applied our algorithm to WGS data from 3,001 ALS patients who have been tested for the presence of the *C9orf72* repeat expansion with repeat-primed PCR (RP-PCR). Taking the RP-PCR calls as the ground truth, our WGS-based method identified pathogenic repeat expansions with 98.1% sensitivity and 99.7% specificity. Further inspection identified that all 11 conflicts were resolved as errors in the original RP-PCR results. Compared against this updated result, ExpansionHunter correctly classified all (212/212) of the expanded samples as either expansions (208) or potential expansions (4). Additionally, 99.9% (2,786/2,789) of the wild type samples were correctly classified as wild type by this method with the remaining two identified as possible expansions. We further applied our algorithm to a set of 144 samples where every sample had one of eight different pathogenic repeat expansions including examples associated with fragile X syndrome, Friedreich’s ataxia and Huntington’s disease and correctly flagged all of the known repeat expansions. Finally, we tested the accuracy of our method for short repeats by comparing our genotypes with results from 860 samples sized using fragment length analysis and determined that our calls were >95% accurate. ExpansionHunter can be used to accurately detect known pathogenic repeat expansions and provides researchers with a tool that can be used to identify new pathogenic repeat expansions.

## Introduction

Variant callers for small variants such as single nucleotide polymorphisms and small insertions or deletions typically require multiple reads to completely span the full length of the non-reference allele (Raczy et al. 2013; DePristo et al. 2011). For variants that deviate significantly from the reference, alternative methods such as *de novo* assembly can be employed if the variant is not highly repetitive (Iqbal et al. 2012; Li 2015; Weisenfeld et al. 2014; Chen et al. 2016). Because high-throughput whole-genome sequencing (WGS) technologies are currently limited to ~150 base pair read lengths, variant-calling methods that rely on reads aligned to the reference are subsequently limited to repeat lengths less than 150 bases (Narzisi and Schatz 2015). Many pathogenic repeat expansions have repeats spanning hundreds to thousands of base pairs (Gatchel and Zoghbi 2005; Kronquist et al. 2008; Dürr et al. 1996; Gijselinck et al. 2016), so it has been assumed that short-read sequencing technologies may not be able to identify pathogenic repeat expansions (Loomis et al. 2013; Ashley 2016).

A recently discovered hexamer (GGCCCC) repeat expansion in the *C9orf72* locus is a major cause of both ALS and frontotemporal dementia (DeJesus-Hernandez et al. 2011a; Renton et al. 2011; Gijselinck et al. 2012). In particular, the pathogenic repeat length (>30 repeats; >180bp) is present in ~10% of all ALS patients including ~40% of familial ALS cases and ~6-8% of sporadic ALS cases in some populations (DeJesus-Hernandez et al. 2011a; Renton et al. 2011; Gijselinck et al. 2012). The most widely used method to detect *C9orf72* repeat expansions is repeat-primed PCR (RP-PCR) together with fragment length analysis (Akimoto et al. 2014). Interpretation of these PCR results can be challenging due to indels in the flanking regions of the repeat, which can lead to both false positives and false negatives (Akimoto et al. 2014). In addition, these PCR techniques do not provide an estimate of the length of the repeat expansions. Southern blotting is the current gold standard for estimating repeat length, but this method is very challenging to set up; requiring a significant amount of input DNA (generally 10 micrograms) and suffering from imprecise size estimates due to somatic heterogeneity (Akimoto et al. 2014; Buchman et al. 2013). As WGS is widely adopted for use in precision medicine initiatives (Ashley 2016; Marx 2015; Ashley 2015) and large scale research projects, a reliable method is needed that can identify the presence or absence of potentially pathogenic repeat expansions in WGS data and also determine their approximate length without additional tests.

Here, we present a method to genotype short tandem repeats (STRs) from PCR-free, WGS data implemented in a software package named ExpansionHunter. This method can determine the approximate size of repeats from just a few units in length up to large, pathogenic expansions that may be significantly longer than the read length. To quantify the performance of this algorithm we first estimate the repeat lengths of two cohorts of ALS patients, all of whom were independently assessed for the presence of the pathogenic *C9orf72* repeat expansion using RP-PCR, and determine the overall sensitivity and specificity of the assay. In addition, we also demonstrate that this method is generally applicable for detecting other repeat expansions by applying it to a set of 144 samples harboring eight other repeat expansions including those that cause fragile X syndrome, Friedreich’s ataxia and Huntington’s disease. We also demonstrated the improved accuracy of this method for genotyping STRs shorter than the read length compared to an existing method (LobSTR) on 860 samples for which the size of the longest repeat allele had been experimentally determined. These analyses show that ExpansionHunter is a comprehensive tool for genotyping both short and long repeats. Thus, it can be used to test for the presence of known pathogenic repeat expansions, and can easily be extended as a general STR caller to identify novel pathogenic expansions in population and pedigree studies.

## Results

We performed paired-end, PCR-free, WGS at an average depth of 45x using Illumina HiSeq 2000 (100 bp reads) and Illumina HiSeq X (150 bp reads) systems on two cohorts of patients with ALS (see Methods). The first cohort of 2,559 patients was used for the initial methods development and the second cohort of 442 patients was used for subsequent validation of the method. All 3,001 samples were tested for presence of the *C9orf72* repeat expansion with RP-PCR (see Methods) for a total of 2,377 wild type and 182 repeat-expanded samples in the first cohort and 416 wild type and 26 expanded samples in the second cohort. A second RP-PCR test using a different primer set, fragment length analysis and Southern blotting was performed on 68 samples from the initial cohort of which 52 had a pathogenic *C9orf72* repeat according to the first RP-PCR (Supplementary Table 2).

To quantify repeat lengths we developed an algorithm that identifies reads that either: 1) fully span the repeat (spanning reads), or 2) include the repeat and the flanking sequence on one side of the repeat (flanking reads), or 3) are fully contained in the repeat (“in-repeat” reads or IRRs) (Figure 1). For repeats shorter than the read length of the sequence data we calculate the repeat length using spanning and flanking reads (Figure 1). To estimate the lengths of repeats that are longer than the read length we identify and count the IRRs. There are three main hurdles associated with using IRRs to accurately identify repeat expansions that exceed read lengths: 1) identifying IRRs comprised of a potentially error-prone repeat motif, 2) identifying regions in the genome where IRR pairs are systematically (and possibly incorrectly) placed by the aligner, and 3) estimating the repeat length based on the total number of IRRs identified. Here, we describe how we solve these problems to accurately identify and characterize expanded repeats.

**Figure 1.**
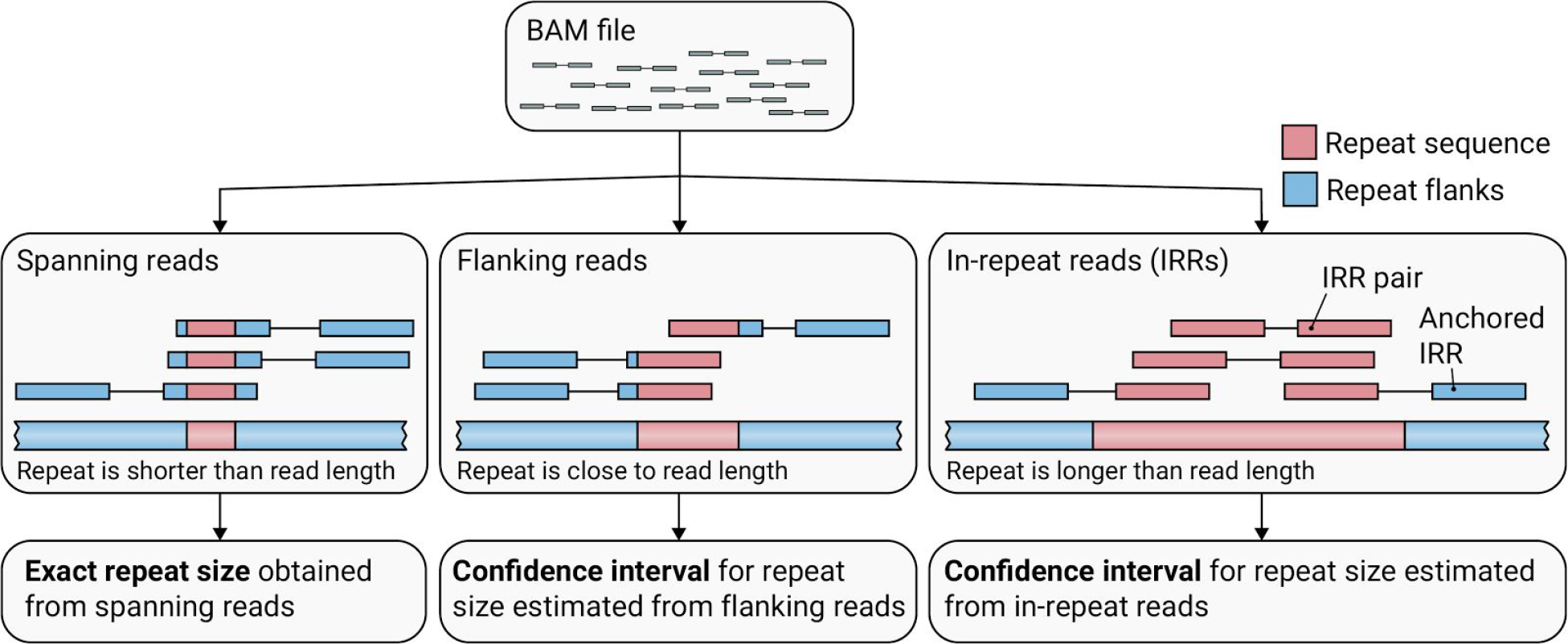
An outline of how ExpansionHunter catalogs reads associated with the repeat locus of interest and estimates repeat lengths starting from a binary alignment/map (BAM) file. (Left) Exact sizes of short repeats are identified from spanning reads that completely contain the repeat sequence. (Middle) When the repeat length is close to the read length, the size of the repeat is approximated from the flanking reads that partially overlap the repeat and one of the repeat flanks. (Right) If the repeat is longer than the read length, its size is estimated from reads completely contained inside the repeat (in-repeat reads). In-repeat reads anchored by their mate to the repeat region are used to estimate the size of the repeat up to the fragment length. When there is no evidence of long repeats with the same repeat unit elsewhere in the genome, pairs of in-repeat reads are additionally used to estimate long (greater-than-read-length) repeats.

### On-target IRRs

Identifying reads originating in highly repetitive regions can be difficult because sequencing error rates are higher in low complexity regions such as homopolymers and STRs (Benjamini and Speed 2012), so we implemented a weighted measure that penalizes base mismatches at low quality bases less than mismatches at high quality bases (see Methods). To identify IRRs that originate within the *C9orf72* repeat we extracted all read pairs where one read is an IRR and the other read aligns with high accuracy (mapping quality (MAPQ) at least 60) within 1 kb of the *C9orf72* repeat locus. We call such reads anchored IRRs. Because the mates of anchored IRRs align to unique sequence near the target repeat we are confident that the IRRs come from the *C9orf72* repeat locus. Anchored IRRs can be used to estimate size of the repeats that are longer than the read length but shorter than the fragment length. For repeats exceeding the fragment length, the number of anchored IRRs provides a lower bound for the repeat length.

### Off-target IRRs

The library preparation used for these sequencing experiments had a mean fragment size of ~350-450 bp but the *C9orf72* repeat expansion can be more than 10 kb in length (Gijselinck et al. 2016). This means that in addition to anchored IRRs, pairs where both mates are IRRs could be present in samples with the *C9orf72* repeat expansion (Figure 1). Because the expanded repeat is not present in the reference, these IRR pairs may not align to the *C9orf72* repeat locus and could either not align at all or misalign to a different locus in the genome (Gijselinck et al. 2016; Church et al. 2015). To identify unaligned or misaligned IRRs, we tested every poorly mapped (MAPQ=0) read in all 182 expanded ALS samples of the first cohort identified by RP-PCR as having the *C9orf72* repeat expansion. These 182 samples contained 29,619 poorly mapped IRR pairs altogether, 33% of these were unaligned and 67% resided in 29 loci (which we term off-target regions), and only 0.1% were located elsewhere (see Methods). Conversely, when we performed the same analysis on 182 random samples without the *C9orf72* repeat expansion according to RP-PCR we did not find IRR pairs in any genomic locus.

We next analyzed positions where the mates of anchored IRRs aligned in all 2,559 samples from cohort one. For each sample we collated the number of anchored IRRs and then grouped IRRs anchored within 500 bp of one another. The *C9orf72* repeat locus had many anchored IRRs in nearly all samples with a pathogenic repeat expansion (178 samples had five or more anchored IRRs and 160 had 10 or more) indicating that the repeat exceeds the read length in these samples as expected. Only 10 genomic loci had more than one IRR anchored outside of the C9orf72 repeat locus in any of these samples (Figure 2). Based on this, we considered all IRR pairs to originate from the *C9orf72* repeat locus and included them in the size estimation when testing this repeat.

**Figure 2.**
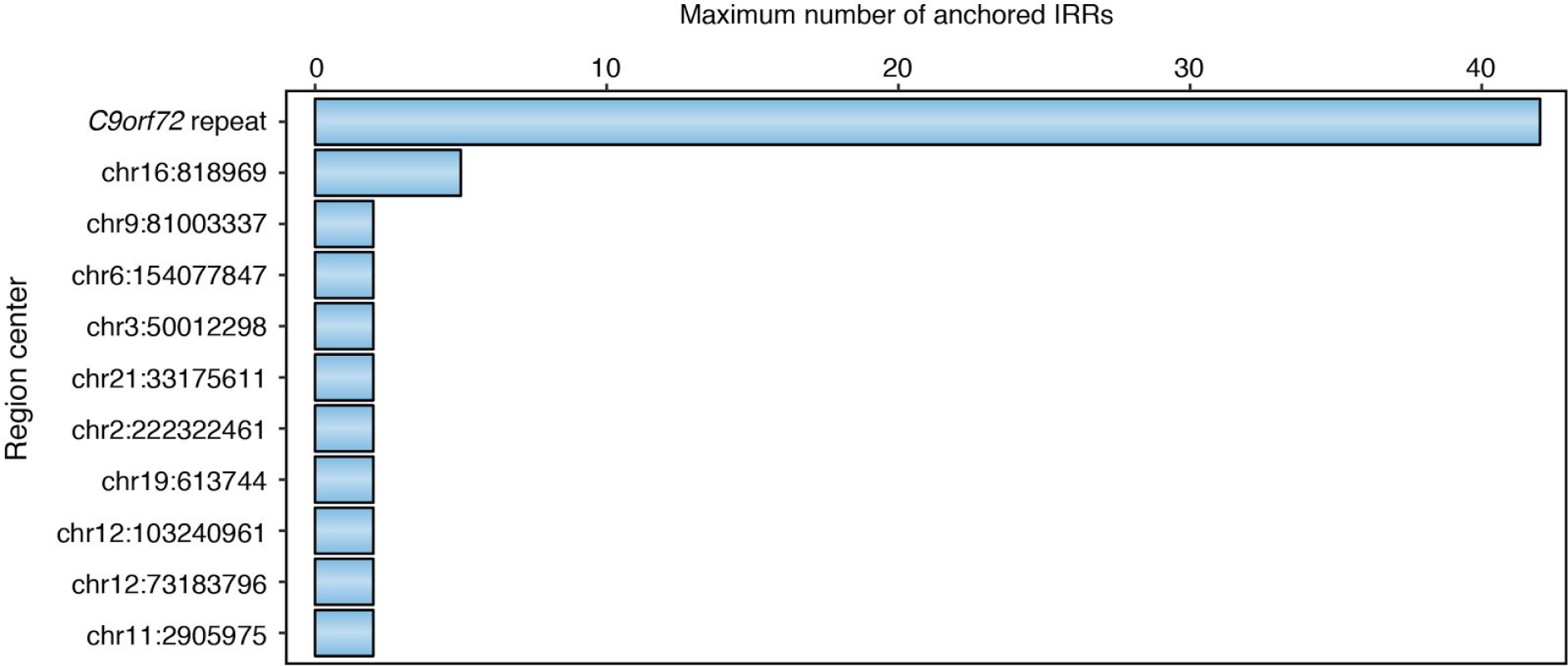
The maximum number of anchored IRRs observed in any of the 2,559 samples from cohort one for the genomic loci with at least two anchored IRRs in at least one sample (se Methods).

### Repeat size estimation

Improvements to short read sequencing technology such as PCR-free sample preparation minimize the GC bias that previously bedeviled PCR-based WGS data (Meienberg et al. 2016). This is illustrated by the improved coverage of high GC regions such as the *FMR1* repeat (Supplementary Figure 5). These improvements enabled us to estimate the length of a region by the number of reads that originate from it even for regions with high GC content. By assuming that the number of reads that originate in a given region follows a binomial distribution we were able to estimate the size of the repeat by the number of IRRs. The number of IRRs in individual samples ranged from 0 to 1,314 corresponding to estimated *C9orf72* repeat sizes of up to 7,152 bp.

For shorter alleles, the sizes of repeats were determined using spanning reads (Figure 1). For repeats that are close to the read length, the repeat may be too long to produce spanning reads but too short to produce IRRs. Therefore, the algorithm also uses flanking reads (Figure 1) to estimate the repeat size (see Methods). In the 2,559 samples of cohort one 1.6% (40) of the samples had a repeat size estimated using only flanking reads that resulted in repeat size estimates from 18 bp to 144 bp (Supplementary Table 2).

ExpansionHunter computes the maximum-likelihood genotype consisting of candidate repeat alleles determined by spanning, flanking, and in-repeat reads (see Methods). When both alleles are longer than the read length, the algorithm computes intervals for possible sizes of short and long repeats based on the two extreme cases: 1) all reads come from one haplotype or 2) half of the reads come from each haplotype.

### Pathogenic *C9orf72* repeat expansion determination

The *C9orf72* repeat sizes for both ALS cohorts were determined with our method and compared to the original RP-PCR results (Supplementary Table 2). Cases where the estimated confidence interval for repeat size overlapped the pathogenic *C9orf72* repeat size cutoff, i.e. the lower bound was less than 30 repeats and the upper bound was greater than 30 repeats, were defined as “grey” and considered “long” in all sensitivity/specificity calculations. Using the RP-PCR calls as the ground truth, the overall sensitivity and specificity for the WGS-based calls were 98.6% and 99.6%, respectively (Table 1). There were 11 samples with a discrepant classification between our method and the RP-PCR in the two cohorts. Eight were “EH positive/RP-PCR negative” (positive=expansion; negative=normal), however, each of these discrepant calls had at least 13 anchored IRRs, which constitutes strong supporting evidence for a pathogenic repeat expansion in these samples (Supplementary Table 4 and Supplementary Figure 6). Predicting the repeat length using only the anchored reads also supported the pathogenic repeat expansion sizing in all eight “EH positive/RP-PCR negative” samples. Conversely, two of the three “EH negative/RP-PCR positive” samples had compelling read-level evidence supporting their negative status: the read-level data supported repeat alleles of two distinct sizes, each spanning fewer than 30 repeat units. Specifically, one sample contained 10 spanning reads with a repeat of size 2 and 10 spanning reads with a repeat of size 5 and the other sample had a size estimate just under the pathogenic cutoff (16 to 26 repeat units). The final “EH negative/RP-PCR positive” sample had just one allele identified (two repeats) but the number of spanning reads (38) was consistent with the read depth (mean depth = 44x) in this sample supporting a homozygous, non-pathogenic variant (Supplementary Table 4 and Supplementary Figure 6).

**Table 1.**
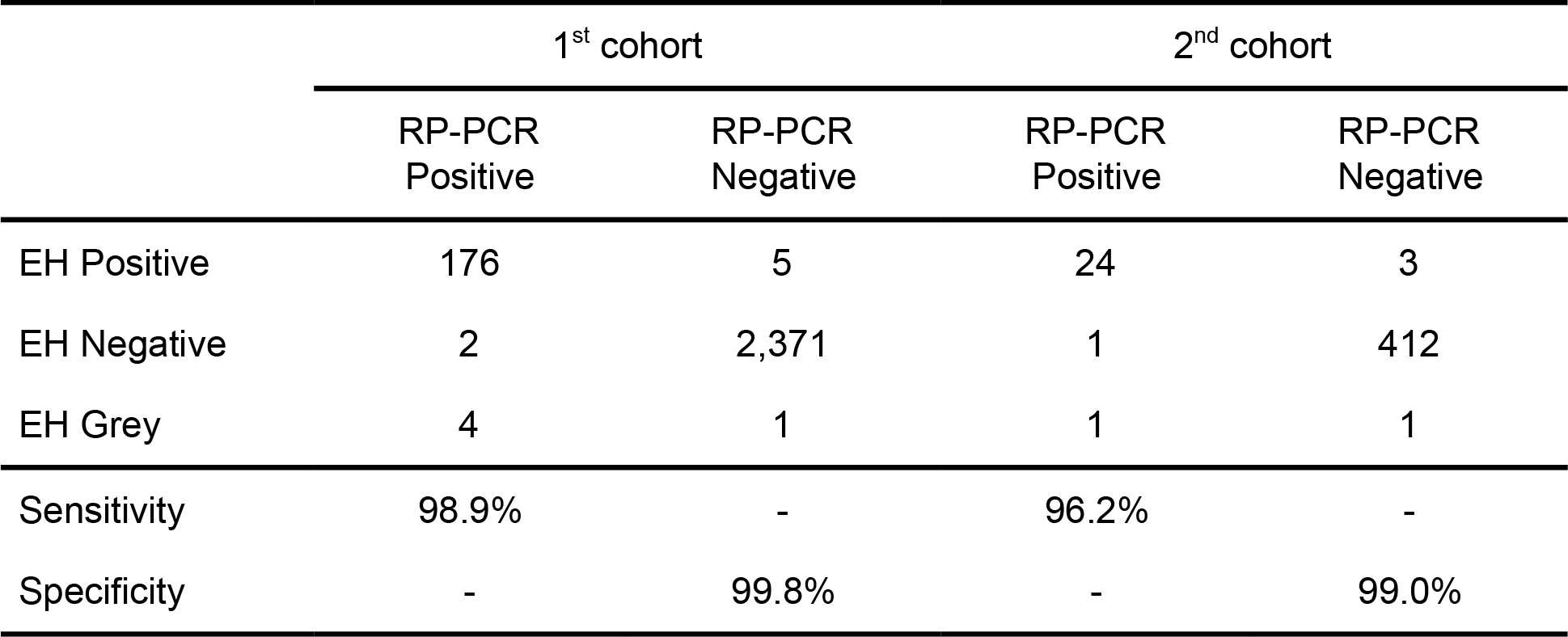
Sensitivity and specificity of *C9orf72* repeat expansion detection by ExpansionHunter (EH) on the ALS samples taking RP-PCR results as the ground truth. EH/RP-PCR Positive (Negative) category refers to samples classified as having expanded (non-expanded) *C9orf72* repeat by each method. EH Grey calls have confidence interval overlapping the pathogenic cutoff (30). Grey calls were considered expanded when calculating sensitivity and specificity.

|  | 1<sup>st</sup> cohort |  | 2<sup>nd</sup> cohort |  |
| --- | --- | --- | --- | --- |
|  | RP-PCR Positive | RP-PCR Negative | RP-PCR Positive | RP-PCR Negative |
| EH Positive | 176 | 5 | 24 | 3 |
| EH Negative | 2 | 2,371 | 1 | 412 |
| EH Grey | 4 | 1 | 1 | 1 |
| Sensitivity | 98.9% | - | 96.2% | - |
| Specificity | - | 99.8% | - | 99.0% |

For the 11 conflicting calls, we reevaluated the original RP-PCR calls and performed an additional RP-PCR and fragment length analysis when our re-assessment of the original RP-PCR call was not conclusive. In 10 of the 11 conflicting calls, we determined that the original RP-PCR call was incorrect and was therefore not conflicting with the ExpansionHunter results (Supplementary Table 4). The remaining conflict produced results that were consistent with the ExpansionHunter call when an additional RP-PCR was performed on this sample with different primers (see Methods). After modifying our calls to incorporate this additional assessment, the only remaining discrepancies in classification are due to the seven “grey” calls where the samples likely have repeat lengths close to 30 repeats. Because we consider “grey” calls as expanded, this method produced just three false positives and no false negatives (EH Grey/RP-PCR Negative).

### Repeats shorter than the read length

To quantify the accuracy of our method for alleles shorter than the read length, we compared our results to those obtained on 860 samples for which the size of the longest allele was estimated using fragment length analysis (Supplementary Table 2). In addition, we also analyzed these samples using the STR calling tool LobSTR (Gymrek et al. 2012). It should be noted that LobSTR is designed for general genome-wide STR calling based on spanning reads and is limited to calling repeat lengths shorter than the read length so it may not make a call for longer repeats. In this comparison, the ExpansionHunter calls agreed with the fragment length analysis in 821 (95.5%) of the samples and the LobSTR calls agreed with the fragment length analysis in 734 (85.3%) of the samples. Of the 39 ExpansionHunter repeat sizes that did not agree with the fragment length analysis, 20 (51%) were in agreement with the LobSTR calls and the remaining 19 calls were predicted to be longer repeats (spanning eight or more repeat units) where LobSTR is less likely to make a call (Supplementary Tables 2 and 5).

Next, we analyzed the 1,770 samples that were sequenced with 2x150 bp reads to get the distribution of the repeat lengths identified from spanning reads in the *C9orf72* repeat. The distribution achieved by this analysis is very similar to the results obtained in a previous study (van der Zee et al. 2013) that used an alternative repeat-primed PCR assay and a short tandem repeat (STR) fragment length assay with flanking primers optimized for alleles with high GC content (STR-PCR) allowing exact sizing of normal lengths (Figure 3). This indicates that we can accurately resolve the length of the short repeats. Because of the requirement for reads to fully span the STR, the maximum repeat size called by LobSTR is 11 repeats even though 4.2% (145 of 3,394) of our alleles are sized greater than 11 repeats. Calling the full spectrum of repeat lengths will enable ExpansionHunter to discover long repeats in population or pedigree studies.

**Figure 3.**
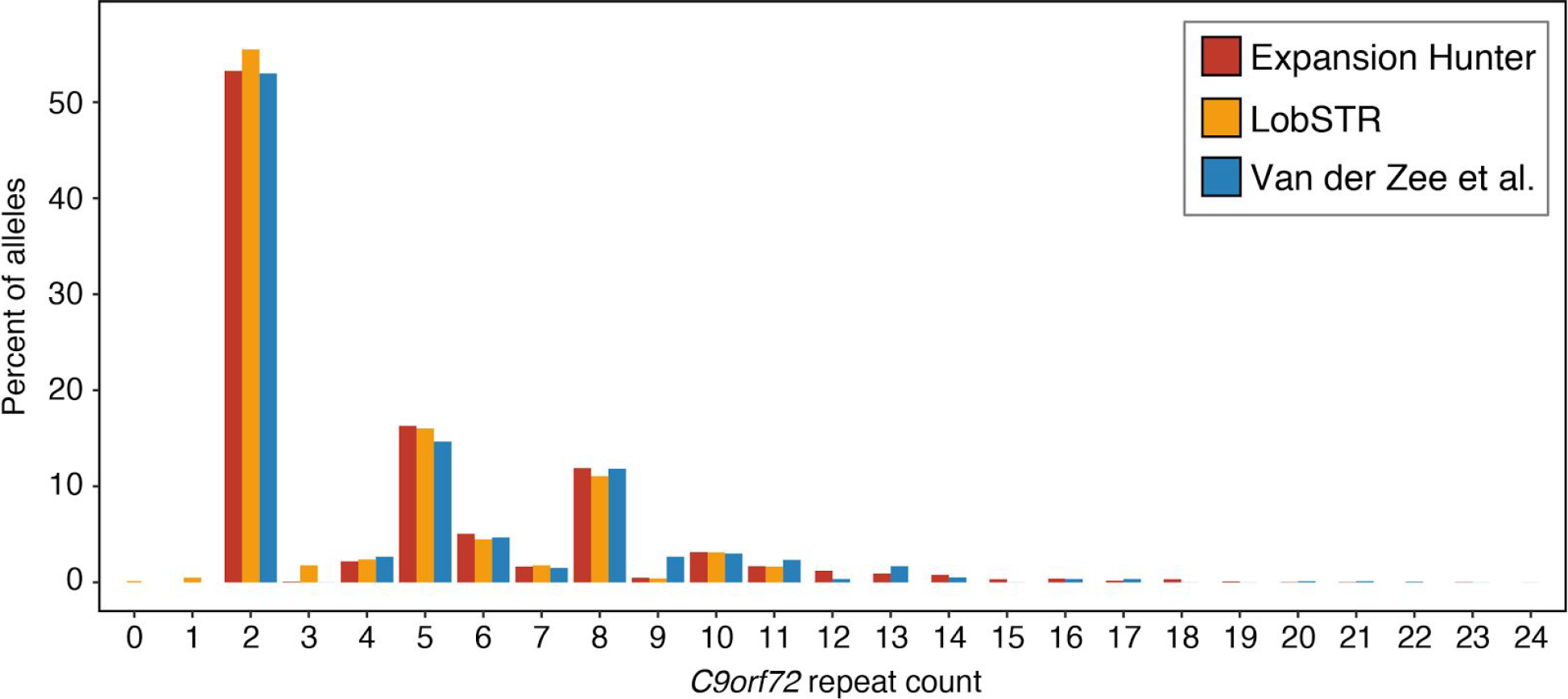
Distribution of EH and LobSTR allele sizes of the *C9orf72* repeat in the 1,770 samples with 150 bp reads from cohorts one and two, compared with those of the FTLD cohort of 318 samples from a previous study (van der Zee et al. 2013).

### Applying ExpansionHunter to other repeat expansions

In addition to the *C9orf72* repeat, several other pathogenic repeat expansions have been identified (McMurray 2010). To demonstrate the general applicability of our method, we tested eight other repeat expansions by sequencing and genotyping 144 samples with known expansions and 25 controls. The sample set contains 90 Coriell samples (https://catalog.coriell.org) with a variety of repeat expansions associated with dentatorubral-pallidoluysian atrophy (DRPLA, *ATN1* gene); fragile X Syndrome (FXS, *FMR1* gene); Friedreich’s ataxia (FRDA, *FXN* gene); Huntington’s disease (HD, *HTT* gene); myotonic dystrophy type 1 (DM1, *DMPK* gene); spinocerebellar ataxia type 1 (SCA1, *ATXN1* gene); spinocerebellar ataxia type 3 (SCA3, *ATXN3* gene); spinal and bulbar muscular atrophy (SBMA, *AR* gene). In addition to the Coriell samples, our data include 54 samples with *HTT* expansions (The U S –Venezuela Collaborative Research Project and Wexler 2004). These 54 samples were processed with a different alignment software (Li and Durbin 2009) confirming that ExpansionHunter is compatible with other commonly-used short read aligners.

Taken together, these 144 samples represent a variety of different repeat expansions with normal/premutation transitions ranging between 87 and 165 bases and premutation/expansion transitions ranging between 114 and 600 bases. The repeats in the *HTT*, *ATXN*1 and *AR* genes are short enough that anchored IRRs alone are sufficient to detect the expansion. For the expansion in the *FMR1* gene, we included off-target reads using the methodology we developed for the *C9orf72* repeat to improve our ability to quantify large repeats. We did not include off-target locations for the other, potentially long repeats because the corresponding motifs (CAG and AAG) are common enough that we could not resolve which repeat the IRR pairs originated from.

Figure 4 depicts the sizes of the longer repeat allele determined by ExpansionHunter. Each of the 144 samples was tested for 8 repeat expansions, one of which is expected to be expanded and the rest wild type. All 24 control samples were similarly tested across all 8 expansions, to assess the relative false positive rate. Our method identified all repeats expected to be pre-mutated (orange circles) or expanded (red circles). The categorization was correct for all repeats with an exception of the *FMR1* repeats where 15 out of 16 repeats were estimated by ExpansionHunter to be pre-mutations instead of full expansions. This discrepancy could be due to mosaicism of expanded *FMR1* repeats (in fact, several of these samples were identified as mosaic in the Coriell database). In addition to mosaicism, other factors such as higher error rates and GC biases may play a role in causing us to underestimate the size of these long repeats.

**Figure 4.**
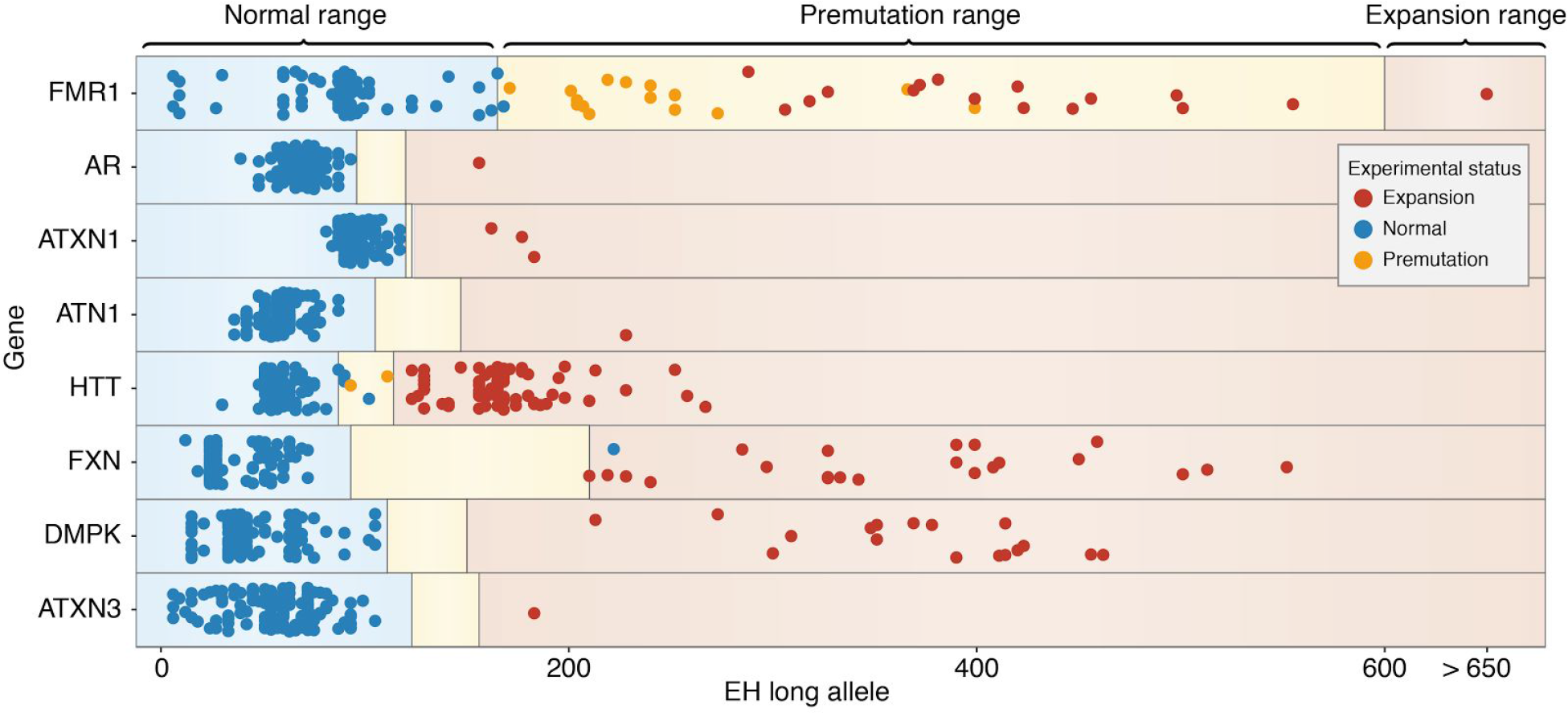
Sizes of the longer repeat alleles predicted by ExpansionHunter in the 144 samples identified as having either a premutation or an expansion at loci associated with eight different diseases and an 24 additional control samples. Circles indicate the most-likely repeat length in bp for a sample identified with a premutation (orange) or expansion (red) and the blue circles show the predicted repeat lengths for the controls. The controls include samples with measurements showing that they fall in the “normal” range and samples that have a different repeat expansion. Thus each sample will have one circle for each of the eight repeat expansions. The regions are shaded to indicate the normal ranges (blue), premutation ranges (yellow) and expansion sizes (light red) (McMurray 2010). Additional information is available in Supplementary Tables 7 and 8.

While we correctly identified all of the expansions, there was one “control” sample showing the *FXN* expansion and two “control” samples with the *FMR1* repeat size at the low end of the premutation range. Both of these results are unsurprising due to the higher carrier frequencies for these two repeats: the carrier frequency is 1:90 for *FXN* (Zamba-Papanicolaou et al. 2009), and 1:178 for the *FMR1* premutation (Hantash et al. 2011). The final three putative FP samples were identified in the *HTT* repeat and include a mother and son who were both sized at 30 repeats (bottom of premutation range) and a third sample with 34 repeats which is small enough for an individual to be unaffected. Visual inspection of the reads supported the ExpansionHunter calls in these samples.

Though ExpansionHunter is designed to work with unbiased (e.g. PCR-free) sequence data, 12 of the samples studied here were sequenced with a PCR step in the sample preparation. These comprised nine samples with either a premutation or expansion at the *HTT* gene and three controls. These samples were correctly classified for the *HTT* repeat despite the high GC content of this CAG repeat (67%). Conversely, the *FMR1* repeat length could not be assessed for all but one of these samples: four samples had no reads covering the repeat and seven were covered very poorly (one to three reads covering the repeat) and produced excessively small repeat lengths. For example, these seven samples were all sized at fewer than 10 repeats whereas for the other 157 samples sequenced without PCR, only four samples had alleles shorter than 20 repeats and the smallest of these was 14 repeats. The poor performance obtained for the *FMR1* repeat shows that some important repeats will be missed or poorly called without PCR-free WGS data.

An important result highlighted in Figures 3 and 4 is that ExpansionHunter is able to size both short and long repeats. This will allow researchers to quantify repeat lengths of all STRs genome-wide to agnostically discover novel pathogenic repeat expansions. To demonstrate this, we took the samples described in Figure 4 and compared repeat sizes in samples with known expansions to the rest of the samples for each repeat. Four of the eight repeat sizes were significantly longer in the affected individuals: *FXN* (One-Sided Mann-Whitney U Test; p=1.3×10^-9^), *FMR1* (p=7.9×10^-11^), *DMPK* (p=4.0×10^-12^) and *HTT* (p=2.5×10^-27^). For this test, we included likely asymptomatic individuals in the control group. For example, the individuals identified by Coriell as having just the *FMR1* premutation or just carrying one expanded allele for the autosomal recessive *FXN* expansion are assigned to the control group. Although this analysis represents an extreme case where the disease is fully penetrant and the samples are perfectly phenotyped, it highlights that it is now possible to agnostically discover new pathogenic variants in pedigree or population studies using short read data.

## Discussion

We have developed a software tool that can identify pathogenic repeat expansions from paired-end, PCR-free WGS data. Comparing against the results obtained with a widely-used wet lab protocol for identifying pathogenic repeat expansions in the *C9orf72* locus, ExpansionHunter was able to correctly classify 208 of the 212 expanded samples and 2,786 of the 2,789 wild type samples. Furthermore, the samples with discordant classifications were all identified as potential expansions (grey) by our method. Potential expansions are identified because there is an uncertainty associated with repeats longer than the read length. In a clinical setting, such calls would trigger a follow up analysis and so all of the expansions were flagged in this analysis.

We also demonstrated that our method generalizes to other repeats by correctly identifying the validated repeats from 144 samples with eight other pathogenic repeat expansions. In total, we examined five repeat motifs (CTG, GAA, CGG, CAG and GGCCCC) at nine different genomic locations and demonstrated that ExpansionHunter can detect repeat expansions in a variety of sequence contexts. It is particularly important that our method works on the very high (100%) GC repeats in *FMR1* (CGG) and *C9orf72* (GGCCCC) genes where both coverage biases and error rates may be elevated. Comparing our size estimates with southern blot experiments indicated that our method may underestimate sizes of some very long repeats, particularly those in the *FMR1* and *AR* genes (Figure 4 and Supplementary Table 7). This underestimation may be caused by the mosaic nature of many repeat expansions, in which case ExpansionHunter will report the average rather than the maximum length. Still, the *FMR1* expansions were generally sized as being larger than the *FMR1* premutation samples, indicating that it may be possible to calibrate our size estimates to account for errors not related to mosaicism. Future work will concentrate on quantifying this behaviour and improving the accuracy of our size estimates for these long repeats.

A major benefit of our tool is that it enables researchers to screen for all known repeat expansions using a single whole-genome sequencing run. As the throughput of WGS increases and the cost decreases, WGS may soon become the basis for frontline tests for repeat expansions and other genetic disorders. Theoretically, long reads can also identify many of the longer repeat expansions (Loomis et al. 2013) but those technologies are still too expensive to be routinely employed for whole genome screening. At the same time, because the substitution and indel error rates in these long reads range from 10 to 30% (Bao and Lan 2017; Sović et al. 2016), it may be difficult to confidently classify the repeat when its size is close to the normal-premutation or premutation-expansion boundary cutoffs unless the samples are sequenced to high depth.

In this study we demonstrated that it is possible to use short read data to confidently identify long, pathogenic repeat expansions and also to accurately determine the size of short, non-pathogenic repeats. Because repeat expansions may expand from generation to generation, pathogenic repeats may show little or no linkage disequilibrium with the surrounding variants. Thus, association studies based solely on SNPs may be blind to these highly polymorphic risk alleles. As association studies based on high-depth WGS data become more widespread, it is now possible to discover new, previously undetected repeat expansions by genotyping them across the population with ExpansionHunter.

In general, our approach is unlikely to work with whole-exome sequence data because: 1) many repeats of interest are not exonic and 2) size estimates for large repeats require assumptions about the average number of reads per base. Some important repeats, like the repeat in *FMR1* gene that causes fragile X syndrome, are 100% GC and are underrepresented in sequence data that includes a PCR step during sample preparation.

In this study, we analyzed nine distinct pathogenic repeat expansions. Future work will focus on extending this method genome-wide to query all possible locations that could harbor a pathogenic expansion. Ultimately, once all known and newly identified pathogenic repeat expansions have been validated, all repeat expansions can be identified from a single PCR-free WGS run.

## Methods

### Whole Genome Sequencing

Paired-end, whole genome sequencing was performed using HiSeq 2000 and HiSeq X instruments. For the 1231 samples sequenced on the HiSeq 2000 instruments 2x100bp reads were generated; for the 1770 samples sequenced on the HiSeq X instruments 2x150bp reads were generated, see Supplementary Table 1. Raw reads were aligned using the Isaac aligner. The quality metrics of these 3,001 samples are described in Supplementary Table 6.

### *C9orf72* PCR

Repeat primed PCR (RP-PCR) was performed on 50-300 ng gDNA with 1x FastStart Mix (Roche), 0.9 M Betaine, 5% DMSO, 1 mM MgCl2, 0.2 mM 7-deaza-dGTP, 0.6-1.3 μM F-primer ([6FAM]AGTCGCTAGAGGCGAAA(GC)), 0.3-1.3 μM R-primer (TACGCATCCCAGTTTGAGACGGGGGCCGGGGCCGGGGCC(GGGG)), 0.6-1.3 μM anchor-primer (TACGCATCCCAGTTTGAGACG) in a total volume of 16-30 μl, with this protocol: 15min 95°C; 2 cycles 1min 94°C, 1min 70°C, 3min 72°C; 3 cycles 1min 94°C, 1min 68°C, 3min 72°C; 4 cycles 1min 94°C, 1min 66°C, 3min 72°C; 5 cycles 1min 94°C, 1min 64°C, 3min 72°C; 6 cycles 1min 94°C, 1min 62°C, 3min 72°C; 7 cycles 1min 94°C, 1min 60°C, 3min 72°C; 8 cycles 1min 94°C, 1min 58°C, 3min 72°C; 5 cycles 1min 94°C, 1min 56°C, 3min 72°C; 10min 72°C. The PCR product was analyzed on an ABI 3730 DNA Analyzer (Applied Biosystems) with PeakScanner software (v1.0). A characteristic stutter pattern was considered evidence of a *C9orf72* repeat expansion. Fluorescent PCR was performed as previously described (DeJesus-Hernandez et al. 2011b).

### Confirmation of *C9orf72* RP-PCR results

The presence of a repeat expansion was determined in a blinded fashion using a 2-step PCR protocol (DeJesus-Hernandez et al. 2011b). In brief, genomic DNA was PCR-amplified with genotyping primers and one fluorescently labeled primer, followed by fragment length analysis with an ABI 3730 DNA analyzer and GeneMapper software (v5). A single PCR fragment could either indicate a homozygous variant or a pathogenic repeat expansion. Subjects with a single PCR fragment were selected for RP-PCR, and PCR products were analyzed with an ABI 3730 DNA Analyzer and GeneMapper software. If the RP-PCR revealed a characteristic stutter pattern, these individuals were screened using Southern blotting techniques, as described previously (DeJesus-Hernandez et al. 2011b). Briefly, a total of 7-10 µg of genomic DNA was digested with XbaI (Promega), and electrophoresed in a 0.8% agarose gel. DNA was then transferred to a positively charged nylon membrane (Roche), cross-linked, and subsequently, hybridized with a digoxigenin (DIG) labeled probe. Expansions were visualized with anti-DIG antibody (Roche) and CDP-Star substrate (Roche) on X-ray film.

### Identifying IRRs

To test if a read fully consists of the repeat motif we compared it to the perfect repeat sequence that was the closest match under the shift and reverse complement operations (e.g. a read originating in a CAG repeat can consist of repetitions of either CAG, AGC, GCA in the forward orientation or CTG, TGC, GCT in the reverse orientation). To do the comparison, we defined the weighted purity (WP) score metric that assigns each matching base a score of 1, each low quality mismatch a score of 0.5 and each high quality mismatch a score of −1. After normalization of the sum of per-base scores for the total read length the WP ranges from −1 to 1. We defined IRRs as reads that achieve WP of 0.9 or above (see supplementary information).

### Identifying off-target regions

IRR pairs originating from expanded short tandem repeats (STRs) may align to other genomic locations especially if the STR is short in the reference genome at the target location. We refer to the loci where IRRs may misalign as off-target regions. Identifying off-target regions enables us to reduce the search for IRRs to a few regions instead of the whole genome. In order to obtain off-target regions for the *C9orf72* repeat we searched through the 182 samples in cohort one that had an expanded repeat according to the RP-PCR to identify all the GGGGCC IRRs. The search was performed through the whole genome for read pairs with a low mapping quality (MAQ=0) and a weighted purity score of at least 0.9. The mapping positions of all identified IRRs were merged if they were closer than 500 bp and the resulting 29 loci that were present in 5 or more samples were designated as off-target regions (Supplementary Figure 4) and were used to find additional reads from the *C9orf72* repeat expansion.

### Repeat size estimation from IRRs

We assume that the probability of observing a read starting at a given base follows the Bernoulli distribution with the probability of success parameter π equal to the ratio of the read depth to the read length. Thus, starting positions of the reads occurring in a given region define a Bernoulli process and the number of reads starting in the region follows a Binomial distribution. If *r* is the read length then one of the terminal bases of any IRR must start at least *N-r* bases away from the flanks of the repeat. The probability of observing *i* such reads is 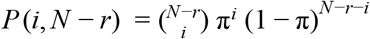. Because we have the estimates for *i* (the number of IRRs) and π (the probability that there is a read starting at a given base), *N* (the repeat size) can be estimated by *r* + *i*/π. The confidence interval for the repeat size is estimated by the parametric bootstrap method (Rice 2007). The same procedure is used to obtain point estimates and confidence intervals for repeat sizes from flanking reads. The confidence interval is truncated according to the size of the longest repeat sequence observed in a flanking read.

### Repeat size determination from spanning reads

The reads spanning the repeat are identified from all the reads that aligned within 1kb of the target repeat region. Each of these reads is tested for the presence of the repeat motif, after which the flanking sequences of the repeat in the read is aligned to the flanking sequences of the repeat in the reference. To be considered spanning, a read must achieve a WP score of 0.9 across the repeat sequence and its flanks. Furthermore, the non-read-length-normalized WP score of the flanking sequences must be at least 2 greater than the score obtained by extending the repeat to the end of the read. In practice, this means that the flanking sequence would have two fewer high quality mismatches compared to extending the repeat or four fewer low quality mismatches. So, if the flanking sequence is similar to the repeat motif then more flanking sequence is required to identify the end of the repeat and the beginning of the flanking sequence.

### Repeat genotyping

Genotype probabilities for repeats of size up to the read length are calculated using a similar model as the one used for SNPs (Li et al. 2009). Namely, *P* (*G* | *R*) = *P* (*R* | *G*) · *P* (*G*) / *P* (*R*) where the genotype G is an n-tuple of repeat sizes and n is the ploidy of the chromosome containing the repeat. The probability *P* (*R* | *G*) is expressed in terms of the probabilities *P* (*ri* | *Hi*) for individual reads *ri* and repeat alleles *Hi* as described in (Li et al. 2009).

Let *k* denote the maximum number of units in a read. For integers 0 ≤ *n*, *m*, *s* ≤ *k* and *p* ∈ (0, 1) we define a frequency function *f*(*m* | *p*, *n*, *s*) ~ *p*(1 − *p*)^*t*^ where *t* = |*n* − *m*| if |*n* − *m*| ≤ *s* and *t* = *s* otherwise. If *r_i_* is a spanning read containing *m* repeat units, *P* (*r_i_* |*Hi* = *n*) = π · *f*(*m* | *p*, *n*, *s*) where π is defined as above (section Repeat size estimation from IRRs). If *r_i_ k* is a flanking or in-repeat read containing *m* repeat units, 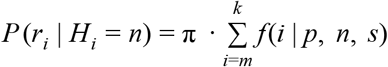. In all our analyses the parameters *p* and *s* were set to 0.97 and 5. The values were chosen to maximize consistency of genotype calls across Platinum Genome pedigree samples (Eberle et al. 2017) on an unrelated set of repeats.

We use read-length-sized repeats as a stand-in for repeats longer than the read length. If only one allele is expanded we estimate the full size of the repeat as described above. If both alleles are expanded, the size intervals are estimated similarly by assuming that between 0 and 50% of in-repeat reads come from the short allele and between 50% and 100% of in-repeat reads come from the long allele.

## Availability

ExpansionHunter is written in C++. The binaries, source code, and documentation are available at https://github.com/Illumina/ExpansionHunter.

## Data access

The raw sequence data used for *C9orf72* repeat expansion detection is stored at SURFsara (Amsterdam) and is available upon request. The raw sequence data for Coriell samples was deposited in the European Genome-phenome Archive (EGA; https://www.ebi.ac.uk/ega) and is available upon request.

The following cell lines/DNA samples were obtained from the NIGMS Human Genetic Cell Repository at the Coriell Institute for Medical Research: NA04724, NA05446, NA0ires39, NA05676, NA06477, NA06591, NA05470, NA05438, CD00014, CD00022, NA03132, NA03200, NA03696, NA03697, NA03756, NA03759, NA03816, NA03986, NA03989, NA03990, NA04025, NA04034, NA04079, NA04567, NA04648, NA04926, NA05131, NA05152, NA05164, NA05185, NA06075, NA06151, NA06889, NA06890, NA06891, NA06892, NA06893, NA06894, NA06895, NA06896, NA06897, NA06903, NA06904, NA06905, NA06906, NA06907, NA06910, NA06926, NA06968, NA07063, NA07174, NA07175, NA07294, NA07536, NA07537, NA07539, NA07540, NA07541, NA07542, NA07543, NA07730, NA07862, NA09145, NA09237, NA09316, NA09317, NA09497, NA13503, NA13504, NA13506, NA13507, NA13508, NA13509, NA13510, NA13511, NA13512, NA13513, NA13514, NA13515, NA13536, NA13537, NA13717, NA14519, NA15847, NA15848, NA15850, NA16197, NA16200, NA16207, NA16209, NA16210, NA16212, NA16213, NA16214, NA16215, NA16216, NA16227, NA16228, NA16229, NA16237, NA16240, NA16243, NA20230, NA20232, NA20233, NA20234, NA20235, NA20236, NA20238, NA20239, NA20240, NA20241, NA20242, NA20243, NA20244, NA23300, NA23374, NA23378, NA23709.

## Acknowledgements

This work was partially carried out on the Dutch national e-infrastructure with the support of SURF Cooperative. This research was supported by NIH/NINDS P01 NS084974 (MvB, RR), R01 NS080882 (MvB, RR), the Thierry Latran Foundation (MAvE, JHV, GPT), the Netherlands Organization for Health Research and Development (Veni scheme, MAvE), the ALS Foundation Netherlands, the MND Association (UK) (Project MinE, www.projectmine.com), and the W. M. Keck Foundation through the grant “Finding Genetic Modifiers As Avenues to Developing New Therapeutics”. Research leading to these results has received funding from the European Community’s Health Seventh Framework Programme (FP7/2007-2013) and Horizon 2020 Programme (H2020-PHC-2014-two-stage; grant agreement number 633413). This study was supported by ZonMW under the framework of E-Rare-2, the ERA Net for Research on Rare Diseases (PYRAMID). This is an EU Joint Programme–Neurodegenerative Disease Research (JPND) project (STRENGTH, SOPHIA, ALS-CarE). The project is supported through the following funding organizations under the aegis of JPND: UK, Medical Research Council (MR/L501529/1) and (ES/L008238/1); Ireland, Health Research Board; Netherlands, ZonMw. AAC receives salary support from the National Institute for Health Research (NIHR) Dementia Biomedical Research Unit at South London and Maudsley NHS Foundation Trust and King’s College London. DEH and CN receives salary from the W.M. Keck Foundation. Samples used in this research were in part obtained from the UK National DNA Bank for MND Research, funded by the MND Association and the Wellcome Trust. We acknowledge sample management undertaken by Biobanking Solutions funded by the Medical Research Council at the Centre for Integrated Genomic Medical Research, University of Manchester.

## References

Akimoto C, Volk AE, van Blitterswijk M, Van den Broeck M, Leblond CS, Lumbroso S, Camu W, Neitzel B, Onodera O, van Rheenen W et al. 2014. A blinded international study on the reliability of genetic testing for GGGGCC-repeat expansions in C9orf72 reveals marked differences in results among 14 laboratories. J Med Genet 51: 419–424.

Ashley EA. 2015. The precision medicine initiative: a new national effort. JAMA 313: 2119–2120.

Ashley EA. 2016. Towards precision medicine. Nat Rev Genet 17: 507–522.

Bao E, Lan L. 2017. HALC: High throughput algorithm for long read error correction. BMC Bioinformatics 18: 204.

Benjamini Y, Speed TP. 2012. Summarizing and correcting the GC content bias in high-throughput sequencing. Nucleic Acids Res 40: e72.

Buchman VL, Cooper-Knock J, Connor-Robson N, Higginbottom A, Kirby J, Razinskaya OD, Ninkina N, Shaw PJ. 2013. Simultaneous and independent detection of C9ORF72 alleles with low and high number of GGGGCC repeats using an optimised protocol of Southern blot hybridisation. Mol Neurodegener 8: 12.

Chen X, Schulz-Trieglaff O, Shaw R, Barnes B, Schlesinger F, Källberg M, Cox AJ, Kruglyak S, Saunders CT. 2016. Manta: rapid detection of structural variants and indels for germline and cancer sequencing applications. Bioinformatics 32: 1220–1222.

Church DM, Schneider VA, Steinberg KM, Schatz MC, Quinlan AR, Chin C-S, Kitts PA, Aken B, Marth GT, Hoffman MM et al. 2015. Extending reference assembly models. Genome Biol 16: 13.

DeJesus-Hernandez M, Mackenzie IR, Boeve BF, Boxer AL, Baker M, Rutherford NJ, Nicholson AM, Finch NA, Flynn H, Adamson J et al. 2011a. Expanded GGGGCC hexanucleotide repeat in noncoding region of C9ORF72 causes chromosome 9p-linked FTD and ALS. Neuron 72: 245–256.

DeJesus-Hernandez M, Mackenzie IR, Boeve BF, Boxer AL, Baker M, Rutherford NJ, Nicholson AM, Finch NA, Flynn H, Adamson J et al. 2011b. Expanded GGGGCC hexanucleotide repeat in noncoding region of C9ORF72 causes chromosome 9p-linked FTD and ALS. Neuron 72: 245–256.

DePristo MA, Banks E, Poplin R, Garimella KV, Maguire JR, Hartl C, Philippakis AA, del Angel G, Rivas MA, Hanna M et al. 2011. A framework for variation discovery and genotyping using next-generation DNA sequencing data. Nat Genet 43: 491–498.

Dürr A, Cossee M, Agid Y, Campuzano V, Mignard C, Penet C, Mandel JL, Brice A, Koenig M. 1996. Clinical and genetic abnormalities in patients with Friedreich’s ataxia. N Engl J Med 335: 1169–1175.

Eberle MA, Fritzilas E, Krusche P, Källberg M, Moore BL, Bekritsky MA, Iqbal Z, Chuang H-Y, Humphray SJ, Halpern AL et al. 2017. A reference data set of 5.4 million phased human variants validated by genetic inheritance from sequencing a three-generation 17-member pedigree. Genome Res 27: 157–164.

Gatchel JR, Zoghbi HY. 2005. Diseases of unstable repeat expansion: mechanisms and common principles. Nat Rev Genet 6: 743–755.

Gijselinck I, Van Langenhove T, van der Zee J, Sleegers K, Philtjens S, Kleinberger G, Janssens J, Bettens K, Van Cauwenberghe C, Pereson S et al. 2012. A C9orf72 promoter repeat expansion in a Flanders-Belgian cohort with disorders of the frontotemporal lobar degeneration-amyotrophic lateral sclerosis spectrum: a gene identification study. Lancet Neurol 11: 54–65.

Gijselinck I, Van Mossevelde S, van der Zee J, Sieben A, Engelborghs S, De Bleecker J, Ivanoiu A, Deryck O, Edbauer D, Zhang M et al. 2016. The C9orf72 repeat size correlates with onset age of disease, DNA methylation and transcriptional downregulation of the promoter. Mol Psychiatry 21: 1112–1124.

Gymrek M, Golan D, Rosset S, Erlich Y. 2012. lobSTR: A short tandem repeat profiler for personal genomes. Genome Res 22: 1154–1162.

Hantash FM, Goos DM, Crossley B, Anderson B, Zhang K, Sun W, Strom CM. 2011. FMR1 premutation carrier frequency in patients undergoing routine population-based carrier screening: insights into the prevalence of fragile X syndrome, fragile X-associated tremor/ataxia syndrome, and fragile X-associated primary ovarian insufficiency in the United States. Genet Med 13: 39–45.

Iqbal Z, Caccamo M, Turner I, Flicek P, McVean G. 2012. De novo assembly and genotyping of variants using colored de Bruijn graphs. Nat Genet 44: 226–232.

Kronquist KE, Sherman SL, Spector EB. 2008. Clinical significance of tri-nucleotide repeats in Fragile X testing: a clarification of American College of Medical Genetics guidelines. Genet Med 10: 845–847.

Li H. 2015. FermiKit: assembly-based variant calling for Illumina resequencing data. Bioinformatics 31: 3694–3696.

Li H, Durbin R. 2009. Fast and accurate short read alignment with Burrows-Wheeler transform. Bioinformatics 25: 1754–1760.

Li R, Li Y, Fang X, Yang H, Wang J, Kristiansen K, Wang J. 2009. SNP detection for massively parallel whole-genome resequencing. Genome Res 19: 1124–1132.

Loomis EW, Eid JS, Peluso P, Yin J, Hickey L, Rank D, McCalmon S, Hagerman RJ, Tassone F, Hagerman PJ. 2013. Sequencing the unsequenceable: expanded CGG-repeat alleles of the fragile X gene. Genome Res 23: 121–128.

Marx V. 2015. The DNA of a nation. Nature 524: 503–505.

McMurray CT. 2010. Mechanisms of trinucleotide repeat instability during human development. Nat Rev Genet 11: 786–799.

Meienberg J, Bruggmann R, Oexle K, Matyas G. 2016. Clinical sequencing: is WGS the better WES? Hum Genet 135: 359–362.

Narzisi G, Schatz MC. 2015. The challenge of small-scale repeats for indel discovery. Front Bioeng Biotechnol 3: 8.

Raczy C, Petrovski R, Saunders CT, Chorny I, Kruglyak S, Margulies EH, Chuang H-Y, Källberg M, Kumar SA, Liao A et al. 2013. Isaac: ultra-fast whole-genome secondary analysis on Illumina sequencing platforms. Bioinformatics 29: 2041–2043.

Renton AE, Majounie E, Waite A, Simón-Sánchez J, Rollinson S, Gibbs JR, Schymick JC, Laaksovirta H, van Swieten JC, Myllykangas L et al. 2011. A hexanucleotide repeat expansion in C9ORF72 is the cause of chromosome 9p21-linked ALS-FTD. Neuron 72: 257–268.

Rice JA. 2007. Mathematical Statistics and Data Analysis.

Sović I, Šikić M, Wilm A, Fenlon SN, Chen S, Nagarajan N. 2016. Fast and sensitive mapping of nanopore sequencing reads with GraphMap. Nat Commun 7: 11307.

The U S –Venezuela Collaborative Research Project, Wexler NS. 2004. Venezuelan kindreds reveal that genetic and environmental factors modulate Huntington’s disease age of onset. Proc Natl Acad Sci U S A 101: 3498–3503.

van der Zee J, Gijselinck I, Dillen L, Van Langenhove T, Theuns J, Engelborghs S, Philtjens S, Vandenbulcke M, Sleegers K, Sieben A et al. 2013. A pan-European study of the C9orf72 repeat associated with FTLD: geographic prevalence, genomic instability, and intermediate repeats. Hum Mutat 34: 363–373.

Weisenfeld NI, Yin S, Sharpe T, Lau B, Hegarty R, Holmes L, Sogoloff B, Tabbaa D, Williams L, Russ C et al. 2014. Comprehensive variation discovery in single human genomes. Nat Genet 46: 1350–1355.

Zamba-Papanicolaou E, Koutsou P, Daiou C, Gaglia E, Georghiou A, Christodoulou K. 2009. High frequency of Friedreich’s ataxia carriers in the Paphos district of Cyprus. Acta Myol 28: 24–26.

